# 3D rotational motion of an endocytic vesicle on a complex microtubule network in a living cell

**DOI:** 10.1101/576702

**Authors:** S. Lee, H. Higuchi

## Abstract

The transport dynamics of endocytic vesicles in a living cell contains essential biomedical information. Although the movement mechanism of a vesicle by motor proteins has been revealed, understanding the precise movement of vesicles on the cytoskeleton in a living cell has been considered challenging, due to the complex 3D network of cytoskeletons. Here, we specify the shape of the 3D interaction between the vesicle and microtubule, based on the theoretically estimated location of the microtubule and the vesicle trajectory data acquired at high spatial and temporal precision. We detected that vesicles showed more frequent direction changes with either in very acute or in obtuse angles than right angles, on similar time scales in a microtubule network. Interestingly, when a vesicle interacted with a relatively longer (> 400 nm) microtubule filament, rotational movement along the axis of the microtubule was frequently observed, implying an obstacle-avoiding motion. Our results are expected to give in-depth insight into understanding the actual 3D interactions between the intracellular molecule and complex cytoskeletal network.

## INTRODUCTION

Extracellular molecules are engulfed by living cells and transported into the intracellular area via the process called endocytosis, which is a crucial pathway for sustaining the life of a cell (1). After the extracellular molecule is internalized into cytoplasmic area, it forms a vesicle surrounded by cell membrane, which is called an endosome (2). How endosomes are transported inside a cell, from the membrane to their destinations, has been one of the major topics in biophysics, in that the movement of endocytic vesicles are not only is coupled with intracellular signaling but also includes key information for biomedical applications, such as drug delivery (3–5).

From the physical point of view, the endocytic vesicles navigate the cytoplasmic area via the interaction with cytoskeletons, such as microtubules and actin filaments, which act as intracellular tracks (6). In detail, a single vesicle is transported as a cargo along the cytoskeletal filament exploiting motor proteins that interact with both the cargo and track (6). The motor proteins are known to selectively mediate the transport: dynein and kinesin are involved in the interaction between the vesicle and microtubule, while myosin V and VI are involved in actin filament-related transport (7–9). Therefore, the research approaches to understanding the mechanism of vesicle trafficking hitherto have primarily focused on revealing the physicochemical properties of a single motor protein (10, 11). These studies increased our knowledge of vesicle movement on the cytoskeleton at the single-molecule scales. Still, many questions remain about the actual movement of vesicles in a living cell, in which the vesicles interact with multiple motor proteins and frequently transfer between multiple cytoskeletal filaments when being transported inside the complex architecture of cytoskeletons (12, 13).

Recent advances in imaging techniques, such as superresolution microscopy, have enabled highly accurate localization of a target vesicles, which has accelerated investigations into how vesicles navigate to their destinations while interacting with multiple cytoskeletons, in the complex geometry of the cytoskeletal network in a living cell (14–16). For example, the vesicle dynamics at the intersection of microtubules was revealed by correlative live-cell and microtubule superresolution imaging (17, 18). In these studies, the behaviors of lysosomes at the microtubule intersection were categorized into pause, pass, and switch-and-reverse, suggesting that the intersection plays a role as an obstacle against the direction of vesicle movement. Additionally, a model for the navigational vesicle that crosses the microtubules was built, and it showed that the separation between microtubules at the crossing influences the vesicle dynamics in a purified environment (19). These studies have helped deepen our knowledge about the actual movement of vesicles in the complex 3D architecture of microtubules.

However, the precise behavior of endocytic vesicles on the complex microtubule network in a living cell, not only at the intersection but also on the cytoskeleton itself, where the active transport occurs, still remains to be investigated. For instance, although it is now known that the vesicle can transfer between the microtubules, the specific 3D shape of movement before and after the transfer is not yet explained. Moreover, 3D movement of vesicles on the microtubule during the active transport in a living cell is still poorly understood in terms of angular and translational motions.

In the present work, we report the detailed features of vesicle movement on a complicated 3D microtubules network structure at high spatiotemporal precision (Fig. S1), achieved by our microscope stage position feedback system (22) that guarantees approximately 3 nm of x, y and 5 nm of axial spatial precision at 100 frames per second. First, the 3D angles between adjacent microtubules where the vesicle sequentially interacted are calculated based on our novel numerical analysis method reported previously (21). Additionally, the relationship between the transfer angle and the time scale is investigated. Second, the rotational motion of a vesicle around the axis of microtubule is detected and investigated in terms of angular and translational velocity. Interestingly, we discovered a characteristic rotational motion of the vesicle with periodically changing angular velocity, which suggests an obstacle-avoidance movement in a long-range transport. Our discovery is expected to inspire further studies on the vesicle transport on the complicated cytoskeletal network in terms of molecular interaction between the vesicle and cytoskeleton.

## MATERIALS AND METHODS

### Cell Culture

The cell line used in the imaging experiment was a GFP-tubulin-labeled KPL4 human breast cell line (23), which was kindly provided by Dr. Kurebayashi (Kawasaki Medical School, Kurashiki, Japan). The cells were cultured in a complete growth medium (Dulbecco’s modified eagle medium with high glucose, Nacalai Tesque) with 10% fetal bovine serum (Thermofisher Scientific, Inc.), L-glutamine (Wako, Inc.), penicillin-streptomycin (Thermo Fisher Scientific, Inc.), and 0.4 mg/ml of G418 (Sigma-Aldrich, Inc.), and incubated at 5% of CO_2_. Endocytic vesicle micropinocytosis was induced by adding 4 nM of carboxyl quantum dots (Qdot 655 ITK Carboxyl Quantum Dots, Thermo Fisher Scientific, Inc.) diluted in PBS to cells, and cells were incubated for 1 hour before performing the imaging. For the imaging experiments, cells were plated on glass-bottom dishes (Matsunami Glass Ind.,Ltd) at a seeding density of 5 × 10^5^ - 8 × 10^5^ cells per dish.

### Live-Cell Imaging

Live-cell imaging was performed via wide-field fluorescence microscopy using an inverted Olympus IX70 microscope equipped with a 60× of oil immersion objective lens (NA 1.4, Plan Apo). The specimen was loaded on the microscope stage and kept at the physiological temperature of 37° C by a heater (Tokai HIT) mounted on the stage. For the stable imaging, a capacitive sensor-based stage stabilization system (22) was attached to achieve high spatial resolution (near 4 nm) in terms of the standard deviation for each x, y, and z coordinate. The dual-focus optics was built by a customized optical path setting, to collect the identical point source at two different focal planes, which were separated by 1 *µ*m in the axial direction. The images from the two focal planes were collected simultaneously by the image sensor of the EMCCD camera (Andor). GFP-tubulin in living cells was excited by a 488 nm laser. The images, filtered by a bandpass filter (500–550 nm, Semrock) were taken for 60 s at 1 fps. Immediately after imaging the GFP-tubulin, the movement trajectory of vesicles labeled with quantum dots was imaged. Quantum dots were excited by a 532 nm laser, and their images were filtered by a 600 nm long pass filter (Edmond), and collected at 100 fps, for > 10 s.

### Super-Resolution Radial Fluctuation (SRRF) Image Processing

For the GFP-tubulin images, post image processing was conducted by Fiji-ImageJ using the plug in ‘NanoJ-SRRF’, which is provided by RHenriques-Lab (20), in order to better visualize the locations of microtubules. The options including ring radius and radiality magnification are all set at the default values.

### Trajectory Data Analysis

The positions of quantum dot labeled vesicles were tracked by customized particle tracking software based on Matlab, which exploited the Gaussian profile of the point spread function. The *x* and *y* values were detected as the center position of Gaussian function, and 𝓏 position was determined by the intensity difference acquired via dual-focus optics (12) and comparison with a predetermined calibration curve. For the vesicle trajectory data, the linear sections were detected where the local curvature of the trajectory was greater than 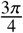, after removing the noise by applying the Gaussian noise filter (window size = 16 data points). The orientation and position of the microtubule for each detected linear section were estimated by using principal component analysis (PCA) (21).

Definitions of various parameters used in the analysis for the detected linear section (21) is shown in Fig. S3. For the position data point set **P**_*l*_ = {P_1_…P_*n*_} in the linear section, **Q**_*l*_ = {Q_1_…Q_*n*_} refers to the corresponding projected point set on the estimated cytoskeleton (red segment). The distance between Q_*α*_ and Q_*β*_, where Q_*α*_, Q_*β*_ ∈ **Q**_*l*_, indicates the pitch length, when there occurs a full rotation. The radius of the rotation is defined as the mean distance between the point set **P**_*l*_ and **Q**_*l*_. The translational movement of the vesicle on the estimated microtubule can be described as ∥Q_*i*+1_-Q_*i*_ ∥at time *t*_*i*_, where *i* refers to the time index. Regarding the direction of rotation, left-handed rotation is defined as when the cumulative angle *ψ*_*i*_ is monotonically decreasing, while the right-handedness indicates a monotonically increasing *ψ*_*i*_. Note that *ψ*_*i*_ = *ρ*_1_ + *ρ*_2_ +… + *ρ*_*i*_, where *ρ*_*i*_ refers to the angle between the vectors 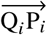 and 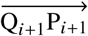. We let 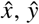 and 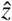 represent the local coordinates based on the right-handed coordinate system, where 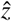 is set to indicate the direction of travel at the local linear section.

### Mean-squared distance (MSD) and mean-squared angular distance (MSAD) calculation

To compare the dynamics of vesicle in the rotational movement section and random movement section, MSD was calculated using following equation.

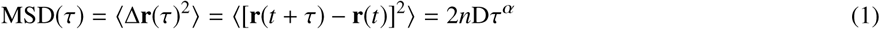

where **r** indicates the three-dimensional position of the vesicle at time *t, τ* refers to the time difference between the position **r** (*t*) and **r** (*t* + *τ*)(24, 25), *n* is the number of dimensions, D denotes the diffusion constant, and *α* is the power determined by the slope of the MSD plot over *τ* (26). Conventionally, active transport of a vesicle is determined when *α* is approximately 2, while passive transport shows *α* ∼ 1 (26, 27). Similarly to MSD, the mean-squared angular displacement can explain the active transport in terms of angle, based on the following equation (28).

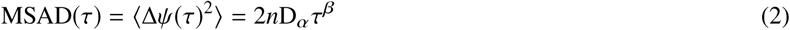

where D_*α*_ indicates the angular diffusion coefficient, *n* is the number of dimensions, *ψ* refers to the rotational angle of vesicle around the estimated axis of cytoskeleton, and *β* is the power determined by the slope of the MSAD plot over *τ*. The calculation results of MSD and MSAD for the detected rotational movement sections and a representative random movement section are shown in Fig. S8.

## RESULTS

### Measurement of 3D angles between microtubules

Conventionally, the angles between microtubule filaments have been measured in two dimensions. However, since microtubules form a three-dimensional network structure, measurement of three dimensional angles considering the geometry of the microtubules in a living cell is required. To measure accurate angles between microtubules in the network structure, the locations of microtubules were estimated from the 3D trajectory of vesicles that apparently interacted with a multiple number of microtubules, based on our numerical method (21). To do this, first, the microtubules labeled by GFP in a living cell were imaged at 1 frame per second (fps) for 60 s (Fig. 1*A*). Immediately after the microtubule imaging, vesicles labeled with quantum dots were imaged for 10 s, at 100 fps. For acquisition of 3D position data of the vesicle trajectory, vesicle-quantum dots were imaged via dual-focus optics (12, 29) within 5 nm of spatial resolution (Fig. S1*D*). The microtubule-GFP images were processed by super-resolution radial fluctuation (SRRF)(20) to accurately define the position of the microtubule, and the trajectories of vesicle-quantum dots were specifically recognized by enhancing S/N by applying Standard Deviation projection to the stacks of vesicle images (ImageJ). By combining the SRRF-processed microtubule image and the vesicle trajectories (Fig. 1*A*), it was possible to determine whether the vesicles were transported along the microtubule.

**Figure 1:**
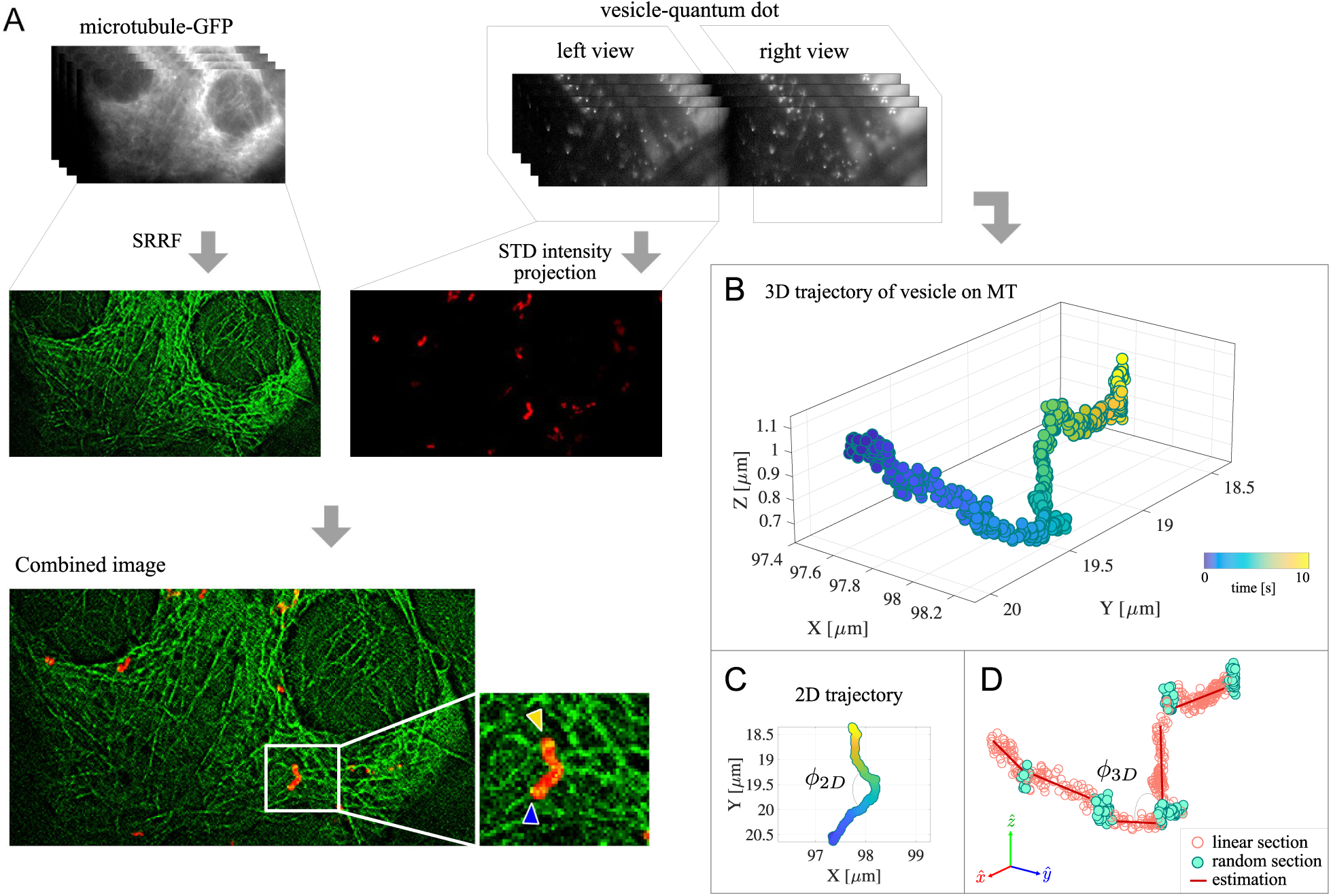
Comparison between the microtubule geometry acquired from 2D and 3D tracking image data. An endocytic vesicle labeled by quantum dots was imaged and tracked in a living cell where the microtubules were labeled by GFP. (*A*) Time-lapse images of microtubules-GFP. Sixty images were taken at 1 fps. Immediately after the microtubule images were taken, vesicle-quantum dot images were taken for 10 s at 100 fps. To acquire specific vesicle trajectories of vesicles that moved on microtubules, SRRF (20) was applied to microtubule images, the standard deviation of the intensity variance of the vesicle image stacks was projected, and the two images were combined (ImageJ). In this combined microtubule-vesicle image, the vesicles moving along microtubules could be detected in two dimensions. (*B*) Three-dimensional trajectory of the vesicle reconstructed by dual focus optics (12). (*C*) *ϕ*_2*D*_ represents the angle that can be roughly estimated from 2D tracking data. (*D*) *ϕ*_3*D*_ indicates the angle between the 3D segments acquired by numerical analysis of vesicle tracking data based on principal component analysis for detected linear sections (21).

As an experimental result, a large portion of all vesicle trajectories were detected on the microtubules (Fig. S2). For a representative trajectory of the vesicle that moved along the microtubules, the 3D reconstruction of the vesicle movement (Fig. 1*B*) indicated that there the vesicle was moved by interacting with multiple numbers of microtubules, while navigating the 3D microtubule network. In the case of a 2D trajectory (Fig. 1*C*), the angle *ϕ*_2*D*_ between 2D microtubule locations can only be roughly assumed, due to the lack of accurate information about the number of microtubules the vesicle interacted with. In contrast, when the vesicle trajectory is reconstructed in three dimensions and analyzed to detect the positions of microtubules (20), now the information about the actual 3D angles between the adjacent microtubules can be clearly obtained (Fig. 1*D*).

### Vesicle transfer between microtubules forms either very acute or obtuse angles

For each trajectory of vesicle that we detected on the microtubule, active transport sections were distinguished as partial linear domains based on the local curvature, and the locations of microtubules were estimated for these linear sections, by using our numerical analysis method (21). A representative vesicle movement on a microtubule was reconstructed in three dimensions, and the data points in the active transport section were colored in pink, while the intermittent pauses or direction changes between active transports were colored in mint green (Fig. 2*A*). For the active transport sections, the estimated locations of microtubules were evaluated using principal component analysis (PCA) (21), and represented by the red-colored segments.

**Figure 2:**
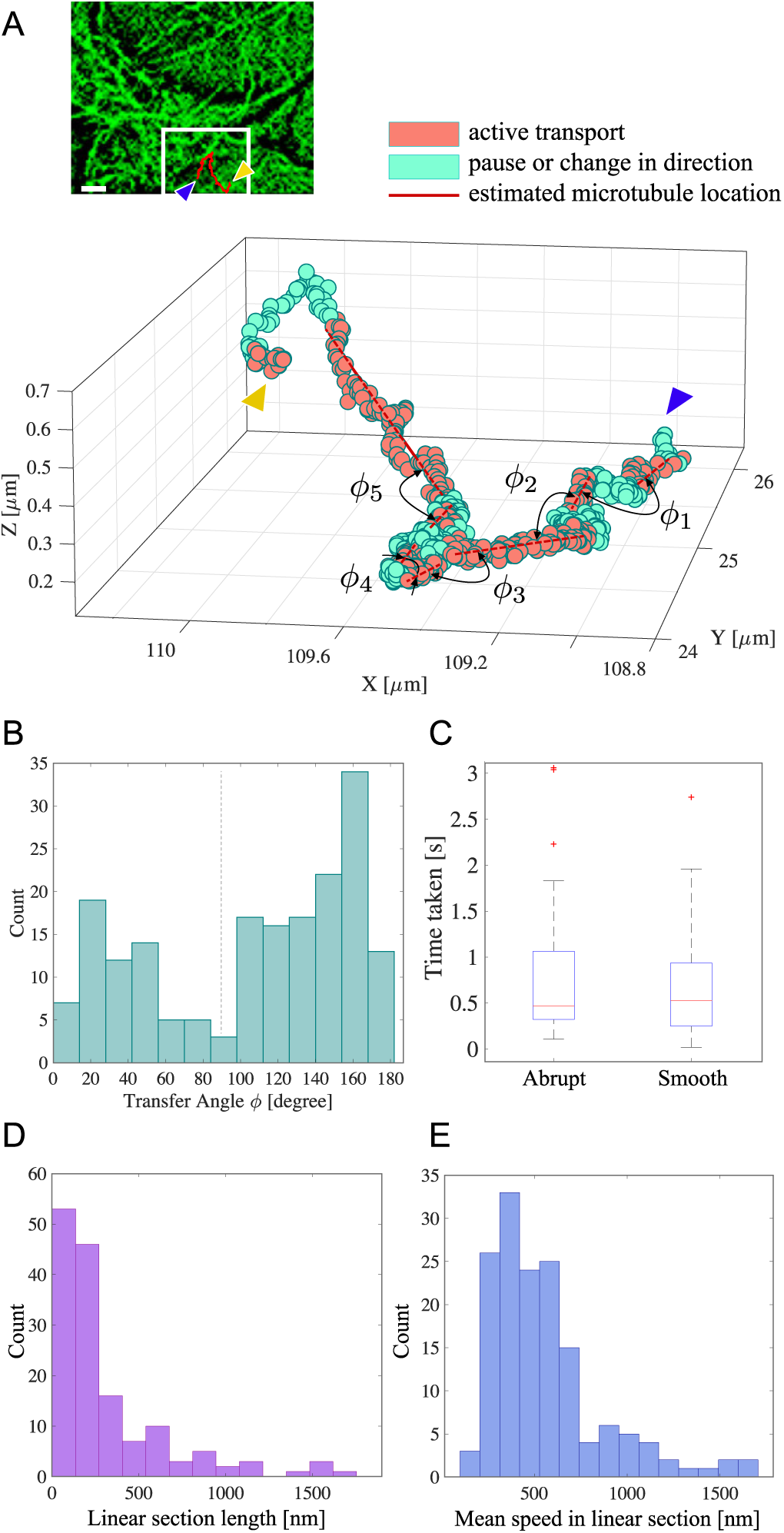
The 3D angles measured between the estimated locations of microtubules. (*A*) Vesicle-quantum dot trajectory (in red) and microtubule-GFP images (in green) were combined to detect the vesicle movement on the microtubule (scale bar = 1 *µ*m), and the 3D trajectory of a vesicle moving in a microtubule network was reconstructed and divided into the linear sections (colored in pink) and pause or direction change sections (colored in mint green) based on the numerical position data analysis reported in our previous work (21). The estimated location of a microtubule was calculated from the linear section via principal component analysis (21). The trajectory started at the point indicated by the blue arrow and finished at the point showed by yellow arrow. The angle *ϕ* was defined as the angle between the adjacent estimated microtubules. (*B*) The distribution of angle *ϕ* was acquired from 61 different trajectories. (*C*) The time taken for the respective case of abrupt direction changes or smooth transfers in a similar direction. (*D*) Distribution of the linear section lengths. (*E*) Distribution of the mean speeds of vesicles in the linear section.

The angle between the estimated location of microtubules was defined as *ϕ*, assuming that the microtubules are approximately linear (21). Since a single trajectory can contain multiple linear sections, as a result of interacting with multiple microtubules, the angles are obtained as *ϕ*_1_, *ϕ*_2_,…, *ϕ*_*n*-1_ for the trajectory which includes *n* estimated microtubules. We acquired 61 different vesicle movement trajectories which included 184 linear sections from the imaging experiment and calculated the angle *ϕ* to investigate the angle at which the vesicle transferred between microtubules. As a result, the angles between microtubules were more frequently measured to be either very acute (10–80°) or obtuse (100–180°), rather than showing angles close to right angles (80–100°). By comparison to the microtubule image (Fig. S2), the transition with an obtuse *ϕ* was either a pause on the same microtubule or a smooth transfer to a different microtubule with a similar orientation. In contrast, an acute *ϕ* indicated an abrupt change in the direction of movement, by switching to a microtubule oriented in the almost opposite direction.

To investigate whether there existed a relationship between the time taken for the transfer and the transfer angle *ϕ*, the time taken for each transfer was measured and compared between the cases of acute angles and obtuse angles. The results indicated that there was not much significant difference in terms of the median time (Fig. 2*C*), as the times in both cases were measured to be approximately 0.5 s. The time shown here can be interpreted as a result of the interaction between the vesicle, motor proteins, and the microtubule at the moment of the transfer. In addition, the lengths of the linear sections were detected as rather short (Fig. 2*D*): 78 % of total linear sections were shorter than 400 nm, with a 176 nm of median value, which implies that the vesicles tended to transfer between microtubules frequently in a microtubule network. The distribution of the mean speeds in the linear sections showed peak values at approximately 500 nm s^−1^ and 1 *µ*m s^−1^ (Fig. 2*E*), which suggest that the vesicles were actively transported via motor proteins in the linear sections.

### Rotational motion of a vesicle with characteristic angular velocity was detected in the linear movement section

Rotational motion around the axis of the microtubule was detected in the linear sections of the vesicle trajectories in the larger data pool, which included the vesicle trajectories that were not always acquired with the microtubule images but were acquired for a longer time span (> 10 s). To investigate the shape of rotation, the quantitative properties of the rotation, such as the rotational angle, handedness, radius, and pitch length, were defined as shown in Fig. S3, based on the notations suggested in our numerical analysis method (21). For a data point P_*i*_ in the linear section, the projection of P_*i*_ onto the position of an estimated microtubule is represented as Q_*i*_, where *i* is the time index. In this case, the right-handed rotation can be defined as when the cumulative angle *ψ*_*i*_ between consecutive vectors 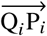 and 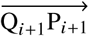, is increasing.

For a representative vesicle trajectory, a left-handed rotational movement was recognized in one of the linear sections (Fig. 3*A*). In this case, the vesicle trajectory was detected around the axis of the estimated microtubule in the anti-clockwise direction, showing monotonically decreasing *ψ*_*i*_ (Fig. 3*B*). We detected not only left-handed rotation, but also right-handed rotation, which showed increasing *ψ*_*i*_ (Fig. 3*C*). For the cases of the complete rotations (*n*=25), there was no significant preference for the direction of rotation, in that 44 % were right-handed rotation and 56 % were left-handed. Additionally, there existed several cases of incomplete rotation with a turning-back motion (Fig. 3*D*). Note that the above three cases, left rotation, right rotation, and turning-back motion, were shown while the vesicles still walked along the axis of the estimated microtubule, with almost constant translational velocity (Fig. 3*B*, *D, E* and Fig. S4). Interestingly, regardless of the direction of rotation, the angular velocities were not constant over time: The angle *ψ*_*i*_ was either monotonically increasing or decreasing with intermittent steady states that were 180° apart (Fig. 3*B*, *C, D*, and Fig. S4). While the angular velocity was uniquely changing, the translational velocity remained constant at the same time. This result implies that there might exist certain tracks of positions on microtubules where vesicles are preferentially carried, and the vesicles quickly dodge around the microtubule when they encounter an obstacle.

**Figure 3:**
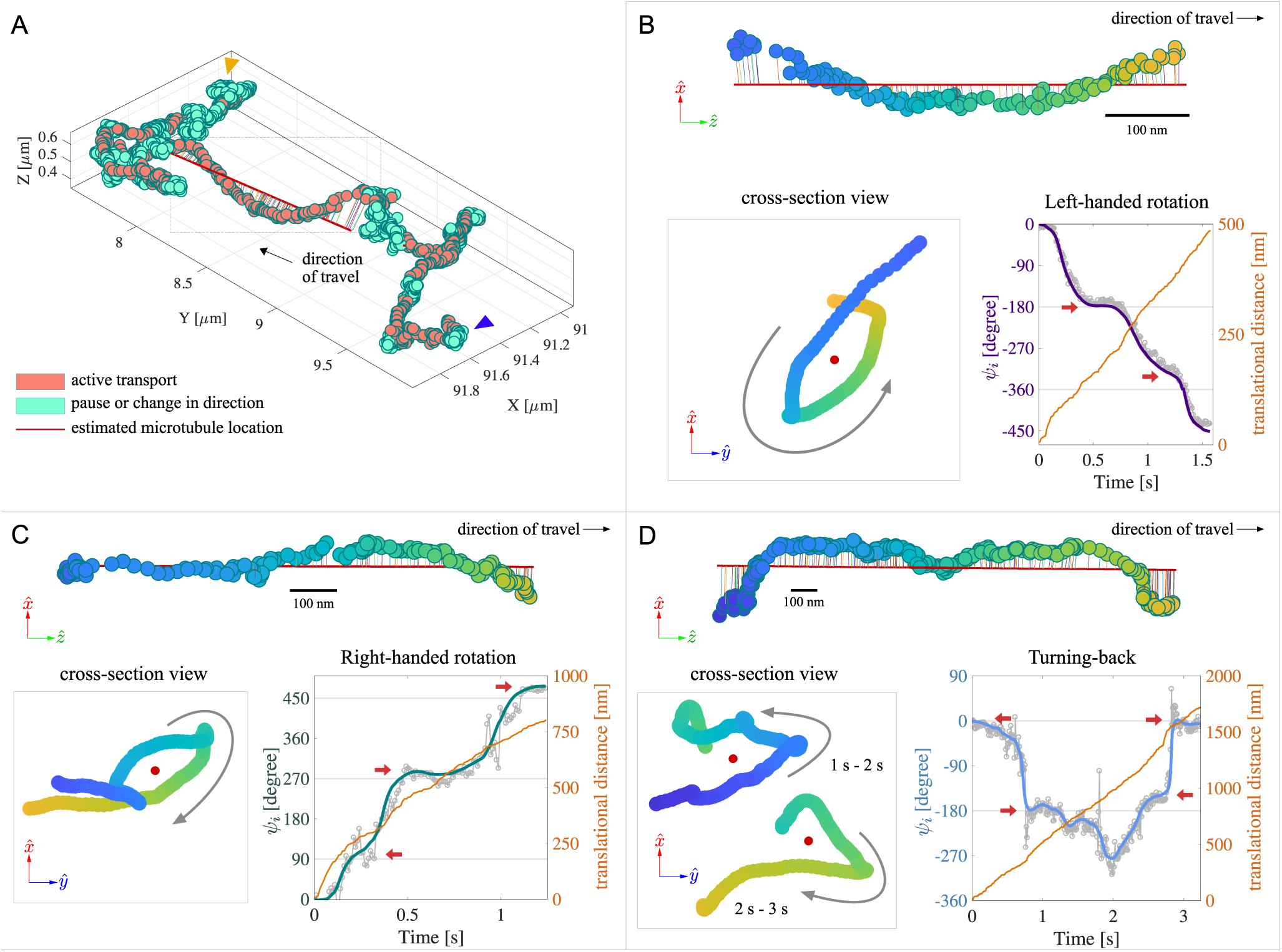
The rotational movement of a vesicle trajectory detected in the linear sections. The data pool of vesicle trajectories has been enlarged by adding the vesicle trajectories that were obtained over a longer time (> 10 s) but were not always taken with the microtubule images. (*A*) A representative trajectory that contains left-handed rotational movement of the vesicle in the linear section. The blue arrow indicates the position where the trajectory starts, while the yellow arrow represents the end point of the trajectory. The linear sections are colored in pink. The linear section showing the rotation along the estimated microtubule segment is indicated in the dashed-line box. (*B*) Left-handed rotation detected in the linear section, shown with a cross-sectional view, and the changes in angular and translational velocities. Cross-sectional view of the rotational movement after the noise is removed by a Gaussian noise filter (see Materials and methods). The red dot in the center indicates the position of the estimated microtubule, which runs inward. The changes in the angle *ψ*_*i*_ over time represents the angular velocity during the rotation. Monotonically decreasing *ψ*_*i*_ represents the left-handed rotational movement of the vesicle around the axis of the estimated microtubule. The small red arrows indicate the inflection points in angular velocity. Note that the inflection points are 180° apart. The changes in the translational distance of the vesicle along the microtubule (colored in yellow) was increased during the rotation, showing almost constant velocity. (*C*) A representative case of right-handed rotational movement of the vesicle, with increasing *ψ*_*i*_ as the vesicle walks along the microtubule. (*D*) A representative case of the incomplete rotation which showed a turning-back motion. Note that the translational velocity is still constant.

### Properties of rotations validate the vesicle movement around the microtubule

To investigate whether these rotational movements were geometrically possible between the vesicle and microtubule, we further examined the pitches, radii, and the translational velocities of the detected complete rotations. We found that the pitches were longer than 400 nm (Fig. 4*A*), occupying > 53% of the probability of occurrence. Compared to the length distribution of all linear sections the median value of which was close to 200 nm (Fig. S5), we can say that the rotation was detected only in the long-range transport as involving a single microtubule filament. The radius of the rotation was defined as the mean distance between the actual position of a vesicle and its projected point on the estimated axis of the microtubule, and the grand mean of the radii was close to 50 nm (Fig. 4*B*), which is plausible considering the sizes of vesicles (radius < 50 nm) and microtubules (radius ∼ 12 nm). The mean translational velocity of the vesicle on the estimated microtubule was measured to be approximately 1 *µ*m s^−1^ (Fig. 4*C*), suggesting that multiple motor proteins, such as dyneins or kinesins, might be involved in the rotational motions.

**Figure 4:**
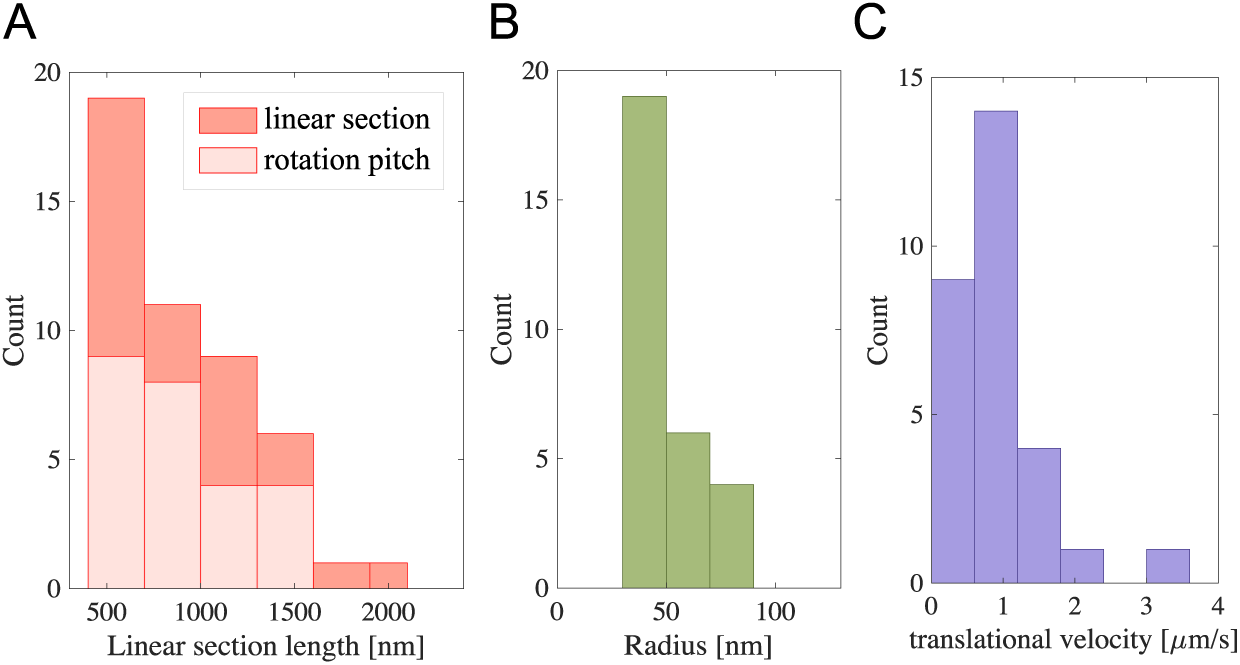
The physical properties of the rotational movement. (*A*) Comparison of length distribution between all linear sections (colored in red) and the pitch lengths in rotations (colored in pink). (*B*) Distribution of the mean radii of the rotational movements (*n*=29). (*C*) Distribution of mean translational velocities on the estimated microtubule. The grand mean value was close to 1 *µ*m s^−1^. The translational velocity refers to the mean of instantaneous speed of the vesicle while walking along the microtubule, mathematically defined as ∥Q_*i*+1_ - Q_*i*_ ∥/*t*_*i*_ where *t*_*i*_ corresponds to the exposure time per frame.

The rotations were not the result of random movement such as Brownian motion but active transport, as it is revealed by the calculation of both mean-squared distance (MSD) and mean-squared angular distance (MSAD) (28, 30) for the detected rotational movements. When the MSD and MSAD for the linear sections which contain rotational movement of vesicles are computed (Material and Method), both MSD and MSAD showed approximately second-order plots over time, implying that the vesicle was actively transported along microtubule during its rotation (Fig. 5 and Fig. S8). For the combined dataset of both incomplete and complete rotations, the mean value of *α* and *β*, the criteria from the power-law fit of MSD and MSAD plots, showed 1.95 and 2.14, respectively (Fig. 5). Since these values are close to 2, it is highly likely that the rotational movements were occurred during active transport of the vesicle (27).

**Figure 5:**
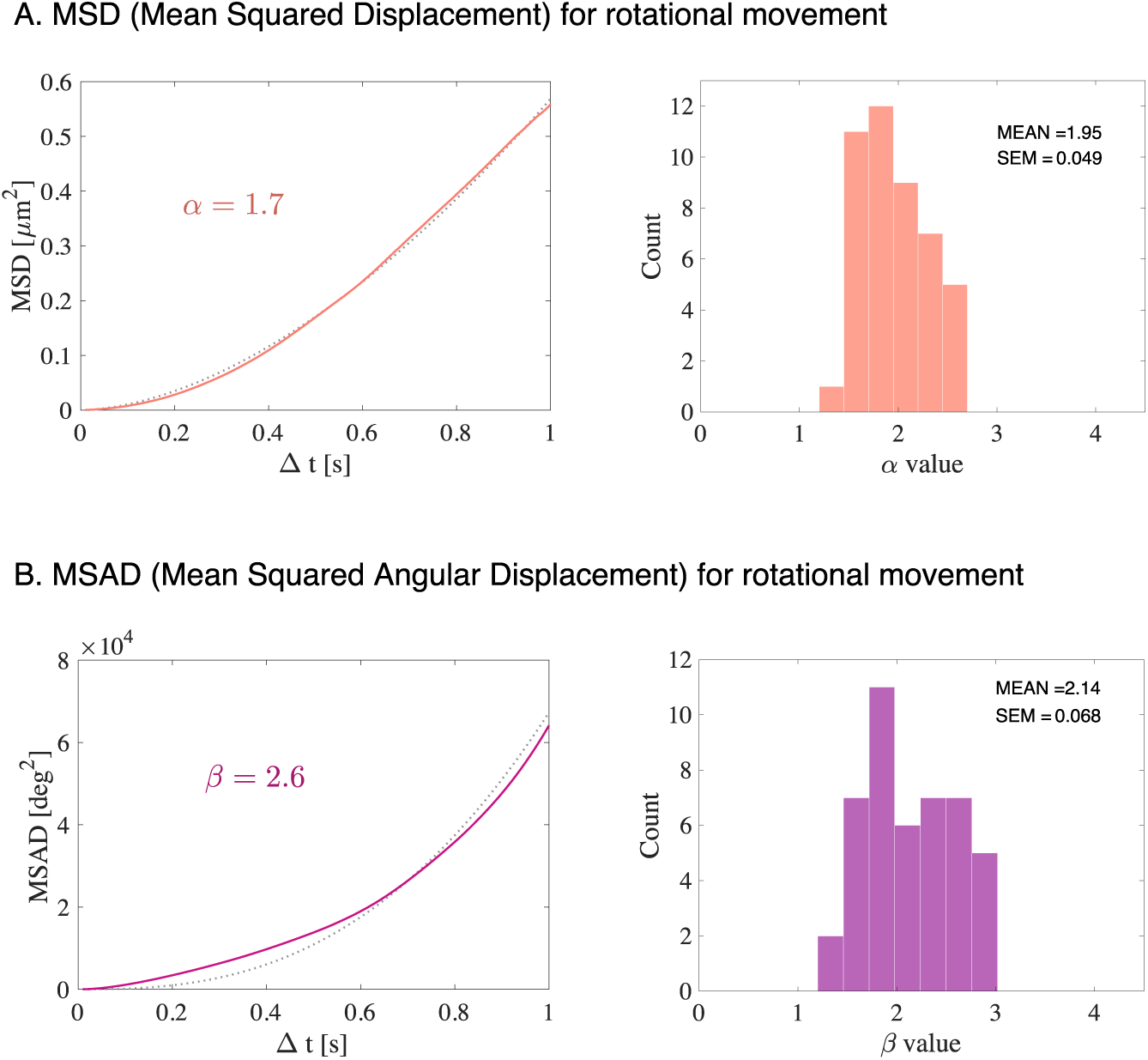
MSD and MSAD for the detected rotational movement. (A) A representative case of *α* calculated from the linear section where rotational movement was observed (left). The mean value of *α* was approximate 1.95 (Right). (B) A representative case of *β* from the rotational movement (left) and the histogram of *β* which produced 2.14 of mean value (Right).

## DISCUSSION AND CONCLUSION

In the presented work, we showed precise vesicle movement trajectory in a complex microtubule network structure in a living cell by analyzing the three-dimensional movement trajectory data of vesicles acquired at a high spatiotemporal precision, achieved by highly stable microscope stage with position feedback system (22). Particularly, we discovered rotational movement of vesicles around the axis of the microtubule, which consists of characteristic quick dodging and slow walking. The precise locations of the microtubules from the movement trajectory of the vesicle are estimated based on our previously reported numerical analysis method (21).

The complex vesicle movement trajectory consists of linear sections and intermittent pauses, which can be understood as a series of the interactions between the vesicle and microtubule (Fig. 2). By computing the three-dimensional angle between the microtubules estimated from the linear sections, we showed that the transfer angle of vesicle was either very acute or obtuse, rarely showing perpendicular direction change (Fig. 2*B*). Compared to the model of angle-independent switching in bead movement at the *de novo* microtubule intersections in cases of obtuse or normal angles (19), our result implies that the probability of orthogonal crosses in the microtubule network in living cells is low, which makes sense considering that the microtubules are basically spread out radially from the microtubule organizing center located near the nucleus. We also measured the distributions of the time taken for the abrupt direction change, the median value of which was approximately 0.5 s (Fig. 2*C*). If the direction change of a vesicle between the microtubules that crossed at an acute angle is caused by the tug-of-war of motor proteins (31–33), the time we evaluated here can be interpreted as the time scale of the tug-of-war between multiple numbers of motor proteins that occurs in the actual 3D cytoskeletal network structure in a living cell condition.

Interestingly, we detected the angular motion of a vesicles that rotated around the axis of the estimated microtubules, especially for longer range transport (> 400 nm). In fact, the rotational movement of a cargo carried by motor protein on the cytoskeleton has been predicted in the previous researches conducted in a purified environment. Dynein showed a bidirectional helical motility with pitch ∼ 600 nm when carrying a vesicle on the microtubule, which was revealed by the bridge assay (34). When we conducted an analysis on the direction of the vesicle transports which showed rotational movements, the results implied that not specific type of motor protein is responsible for both right-handed and left-handed rotational movement (Fig. S6). This is one of the reasons we can consider that the rotational movement is caused by any obstacle the vesicle encountered during the transport.

Additionally, the three-dimensional obstacle-circumventing motion of a vesicle along microtubules was recently reported (18), although the rotations were not found. In the latter study, the circumventing motion of a vesicle was detected when the vesicle encountered an obstacle, such as an intersection of microtubules. Therefore, it is expected the rotational motion detected in our analysis seems to be induced by the obstacles. Our research presents the initial observation and analysis of the actual rotational movement of a vesicle on a microtubule in a living cell, and thus provides insight into understanding the detailed features of vesicle motion on the cytoskeleton, which is a result of the interaction between a vesicle and a multiple number of motor proteins (35).

Regarding the rotational motion of the vesicle around the axis of a microtubule, there were several characteristic aspects: (1) There was no significant preference between left-handed and right-handed rotation. (2) Turning-back motion, without a complete turn, existed. (3) The pitch of rotation was not uniform but rather long (∼600 nm) compared to the length distribution of all the linear sections. (4) The angular velocity was not constant, while the translational velocity was constant, and there existed quick dodging motions between 180°-apart angular positions. If the rotational movement is caused by inherent physical properties of the microtubule, such as its structure, the pitch of rotation should be uniform, rather than distributed broadly. Therefore, considering the properties of the rotation stated above, it is probable that the rotational motion occurs when a vesicle avoids any obstacle while being transported (18). There can be various roadblocks during vesicle transport, which disrupt the uniform and steady interaction between the vesicle and cytoskeletons, as the cytoplasmic area is highly crowded with macromolecules (36). More directly, the intersection of microtubules (17–19) or the existence of microtubule-associated proteins such as tau (9, 35), are some of the obstacles that a vesicle can encounter.

Additionally, there is a very low probability that the detected rotational movement is the result of the vibration or fluctuation of microtubules. This is because the cytoplasm acts as a strong damper, in contrast to many physical models, which have predicted a high frequency (∼ GHz) of microtubule vibration under ideal conditions (37). As a simple experiment, the trajectory of three independent vesicles located on the same microtubule were investigated, and they did not show any synchronized movements in the axial direction (Fig. S7). The other possible influences on the detected rotations, such as the diffusion of vesicles, spin of vesicles, or detection noise, can be ruled out because the characteristic rotational motions were found in 53 % of the long-range transports (> 400 nm), which is unlikely to be caused by random motions. Moreover, since the mean of *α* and *β*, which are the criteria of active transport computed from respective MSD and MSAD plots with power-law fit, showed the values close to 2 (Fig. 5 and Fig. S8). This result reflects that the rotational movements were not random motions but active transports.

Since our result is the initial analysis of 3D rotational movement of a vesicle in a complex cytoskeletal network in a living cell, it is expected that related studies can yield new insight into understanding and analyzing the actual movement of vesicles navigating in the intracellular area.

## AUTHOR CONTRIBUTIONS

S. Lee and H. Higuchi designed research; S. Lee performed experiment and analysis; and S. Lee and H. Higuchi wrote the paper.

## ACKNOWLEDGMENTS

We thank Dr. H. Kim in the University of Tokyo for helpful discussions and critical reading of the manuscript. This work has been supported by Grands-in-Aid for Scientific Research (A and B) (H. H. 23247022 and 16H04773) and Challenging Exploratory Research (H. H. 17K19343) from the Japan MEXT.

